# Osterix Facilitates Osteocytic Communication by Targeting Connexin43

**DOI:** 10.1101/2024.09.09.611984

**Authors:** Zuping Wu, Qian Chen, Qian Gao, Muchun Liang, Yumeng Zhou, Li Zhu, Jiahe Wang, Yang Shen, Junjun Jing, Jing Xie, Xiaoheng Liu, Shujuan Zou, Demao Zhang, Chenchen Zhou

## Abstract

Osteocytes, terminal-differentiated cells in bone, are now considered as more pivotal regulators of mature bone homeostasis than other bone cells, since they constitute 90- 95% of the bone cell population. Given their non-migratory nature within the mineralized matrix, their unique dendrites are crucial for cell-to-cell communication in response to both intracellular and extracellular stimuli, such as bone fracture or mechanical load. Here, we showed that Osterix (Osx), usually recognized as a specific doorkeeper for osteoblast differentiation during new bone formation marked by collagen type I α 1 (Col1α1), was unexpectedly co-expressed with Col1α1 in osteocytes within the cortical bone of mice. Deleting Osx in Col1α1-positive osteocytes disrupted cortical bone structure and osteocytic dendrites in mice, thus impairing transcellular fluid flow and intercellular communication. Conversely, overexpression of Osx in osteocytes enhanced these processes. Furthermore, we identified Connexin43, a critical protein of gap junction channel, was a direct transcriptional target of Osx in regulating dendrites of osteocytes. Pharmacological restoration of Connexin43 levels rescued the dysfunction in Osx-deficient osteocytes both in vitro and in vivo. Taken together, this work demonstrated Osx’s distinct role in osteocyte function through maintaining intercellular signaling, which broadened the current understanding of its role in Col1α1-positive bone cells, extending beyond osteoblasts and bone mineralization, offering new insights into bone diseases such as fracture nonunion or disuse osteoporosis.

## Introduction

The integrity of skeletal health is contingent upon the function of bone cells, with osteocytes playing a more pivotal role in mature bone health than other bone cells, like osteoblasts or osteoclasts. Because osteocytes are embedded within the mineralized bone matrix upon maturation, and approximately tenfold more numerous than osteoblasts, constituting about 90-95% of the total bone tissue cell population(Camal Ruggieri et al., 2021; Dallas et al., 2013). Their star-shaped morphology, characterized by an extensive network of slender dendrites, is integral to their physiological function, given their non-migratory nature within the bone matrix(Tu et al., 2015). So, the dendritic network is crucial for osteocyte intercellular communication, and which is essential for bone remodeling triggered by osteocytes in response to environmental stimuli and signal transduction, regulating osteolysis through osteoclasts and osteogenesis through osteoblasts(Robling & Bonewald, 2020; Tu et al., 2015). Gap junction communication, facilitated by connexin channels, is critical for bone cells to perform their functions(Ma et al., 2019). Connexin43 (Cx43) deficiency in osteoblasts and osteocytes results in fracture nonunion. And aging is associated with a reduction in osteocytic dendrites and a degeneration of the osteocyte network, potentially leading to diminished bone anabolism and a reduced response to mechanical loading.(Tiede-Lewis et al., 2017) Consequently, senescent osteocytes exhibit impaired mechanical sensitivity and an accumulation of cellular senescence(Hayashi et al., 2019; Schurman et al., 2021). Mechanical loads applied to dendritic processes or cell bodies trigger the opening of hemichannels on the osteocytic body. But the hemichannel activity in dendritic processes is not detected when the osteocytic body is mechanically stimulated, indicating that the dendritic processes are key structures for sensing and conducting mechanical stress and its subsequent signal transduction.(Burra et al., 2010; Chen et al., 2019) The dendritc network of osteocytes are in charge of transcellular fluid flow between osteocytes. Collectively, osteocytes play a pivotal role in bone remodeling, with their unique dendrites being essential for cell-to-cell signaling. Yet, many questions remain to be answered about this process.

Osterix (Osx) is known as a master transcription factor that initiates the cascade leading to osteogenesis—the differentiation of osteoblasts into osteocytes, involving in new bone formation.(Hojo et al., 2016; Zhou et al., 2010) Therefore, Osx has been usually recognized as an osteoblast-specific marker of early osteogenesis. Further research has revealed that Osx variants are linked to bone mineral density in both children and adults,(Timpson et al., 2009) suggesting an indirect role for Osx in osteocyte maturation and bone mineral density. However, in this study, we made an unexpected discovery of strong Osx expression within osteocytes, suggesting a direct regulatory role of Osx in osteocyte function. Given the critical role of osteocytes in maintaining mature bone health, it is imperative to elucidate the direct effect of Osx on osteocyte function. In addition, type I collagen is also known as a classical marker for osteoblasts, and Col1α1-CreER of transgenic mice is commonly used to label osteoblasts, whereas we identified that Col1α1-CreER also obviously labeled osteocytes, which broke the previous understanding of Col1α1-positive bone cells. In this work, we utilized a Cre-LoxP system to conditionally knockout Osx and focused on investigating its function in Col1α1-positive osteocytes. Our findings demonstrated that Osx directly supported the osteocyte dendritic network through targeting Cx43, thereby maintaining intercellular communication among osteocytes.

In sum, while previous studies have primarily attributed the function of Osx in Col1α1- CreER-labeled cells to osteoblasts, this work sheds new light on the role of Osx in Col1α1-positive osteocytes, contributing to intercellular signaling. This work broadened our understanding of Osx’s role in maintaining bone homeostasis, offering new insights into bone diseases involving osteocytes, such as fracture nonunion or disuse osteoporosis.

## Results

### Osx obviously expressed in osteocytes while Col1α1-CreER labeled osteocytes

To elucidate the role of Osx in osteocytes, we initiated this study by examining Osx expression within these cells. Immunofluorescence staining revealed that Osx is obviously expressed in osteocytes of the femoral cortical bone in mice (Fig. 1A & Fig. 2A). Meanwhile, utilizing the Col1α1-CreER mouse crossed with tdTomato reporter mice (ROSA26-loxP-stop-loxP-tdTomato), we traced the in vivo expression of Col1α1 and intriguingly found it to also label osteocytes (Fig. 1A). Notably, the expression of Osx (green) and Col1α1 (red) overlaped in osteocytes, as indicated by the yellow merged signal (Fig. 1A), suggesting that Osx knockout using Col1α1-CreER could directly influence osteocyte function in mice. To investigate this, we generated Osx^flox/flox^;Col1α1-CreER conditional knockout (Osx^cKO^) mice, focusing on the role of Osx in osteocytes (Fig. 1B). Successful breeding of cKO mice was confirmed through genotyping (Fig. 1B, Fig. 2A&B, and Fig. 3A&B). In vitro, we modulated Osx expression in osteocytes by knockdown and also overexpression respectively, as validated by Western blots (Fig. 1C-E). Collectively, these findings indicated that Osx was expressed within osteocytes and may play a pivotal role in their function.

**Figure 1.**
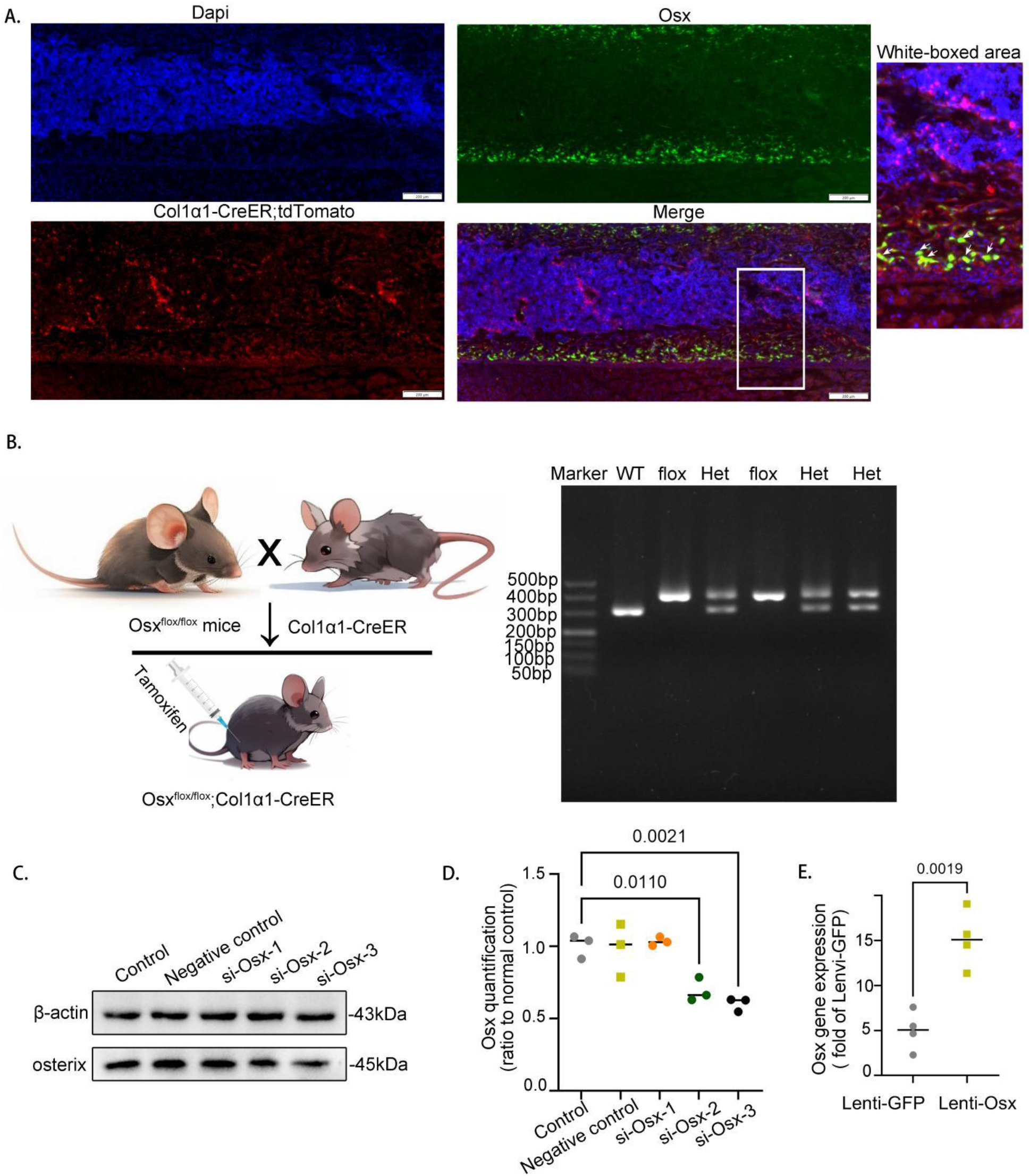
Osx obviously expressed in osteocytes while Col1α1-creER labeled osteocytes. (A) Immunofluorescence staining demonstrated the co-expression of Col1α1 and Osx in osteocytes of mouse cortical bone. Col1α1 = Collagen I α 1. Scale bar, 200 μM. (B) The breeding scheme and genotyping for Osx conditional knockout mice. Het, heterozygote. (C and D) Western blots showed the expression of Osx was repressed by small interfering RNA (si-Osx). The data represented the mean ± SD. Protein levels were normalized to β-actin. *p < 0.05. (E) qPCR showed the mRNA expression of Osx in osteocyte was significantly increased after being transfected with lenti-Osx. Expression levels were normalized using GAPDH and lenti-GFP as controls. *p < 0.05.

**Figure 2.**
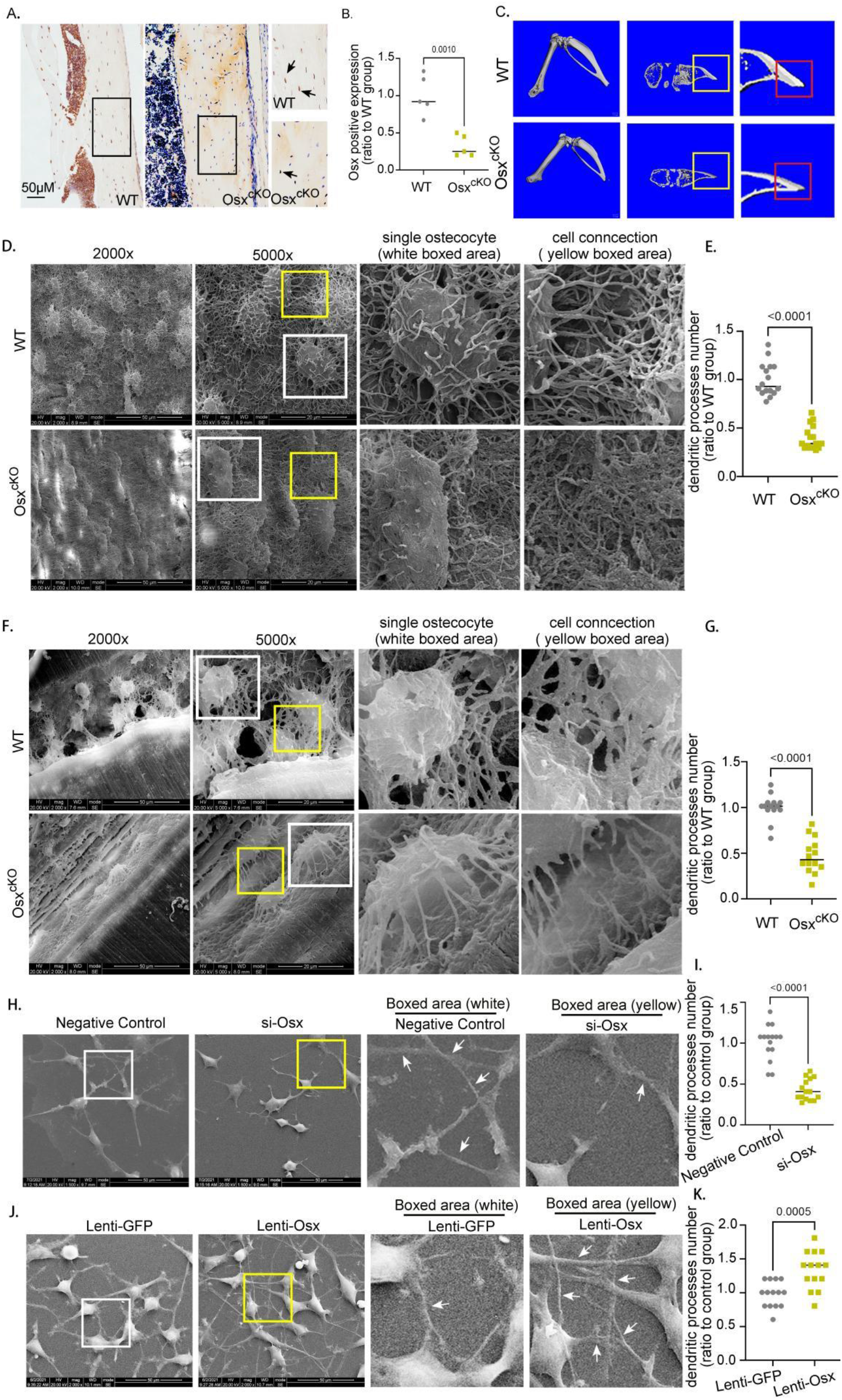
Osx deficiency impaired cortical bone structure and decreased osteocytic dendrites both in vivo and in vitro. (A) Immunohistochemical staining showed a decreased expression of Osx in osteocytes of the cortical bone in Osx conditional knockout (Osx^cKO^) mice when compared to wild-type (WT) group. Scale bar, 50 μM. (B) The reduced expression of Osx in Osx^cKO^ mice was confirmed via quantitative analysis as depicted in (A). The results were presented as the mean ± standard deviation (SD). The data were derived from multiple independent biological samples. *p < 0.05. (C) Representative micro-computed tomography (μCT) images highlighted the differences in tibial cortical bone structure between Osx^cKO^ mice and WT mice. (D and E) Scanning electron microscopy (SEM) images demonstrated a reduction in osteocytic dendrites in the femurs of Osx^cKO^ mice compared to WT mice. The white box delineated the dendritic changes of an individual osteocyte, while the red box showed alterations in intercellular dendrites among osteocytes. The number of dendrites was quantified per osteocyte, with a total of 16 cells analyzed per group. Scale bar, 50 μM & 20μM. *p < 0.05. (F and G) SEM images revealed a similar reduction in osteocytic dendrites in the cranium bone of Osx^cKO^ mice compared to WT mice. The white and red boxes served the same purpose as described in (D&E), and the same quantification method was applied. Scale bar, 50 μM & 20μM. (H and I) SEM images showed a decrease in intercellular dendritic connections of osteocytes in the si-Osx group, where Osx expression was suppressed by si-Osx, compared to the control group. The number of dendrites was calculated per cell, with 16 cells analyzed each group. Scale bar, 50 μM & 20μM. *p < 0.05. (J and K) SEM images demonstrated an increase in intercellular dendritic connections of osteocytes in the Lenti-Osx group, where Osx expression was over-expressed using a lentiviral vector, compared to the control group. The quantification of dendrites was performed per cell, with 16 cells analyzed per group. *p < 0.05. Scale bar, 50 μM & 20μM.

**Figure 3.**
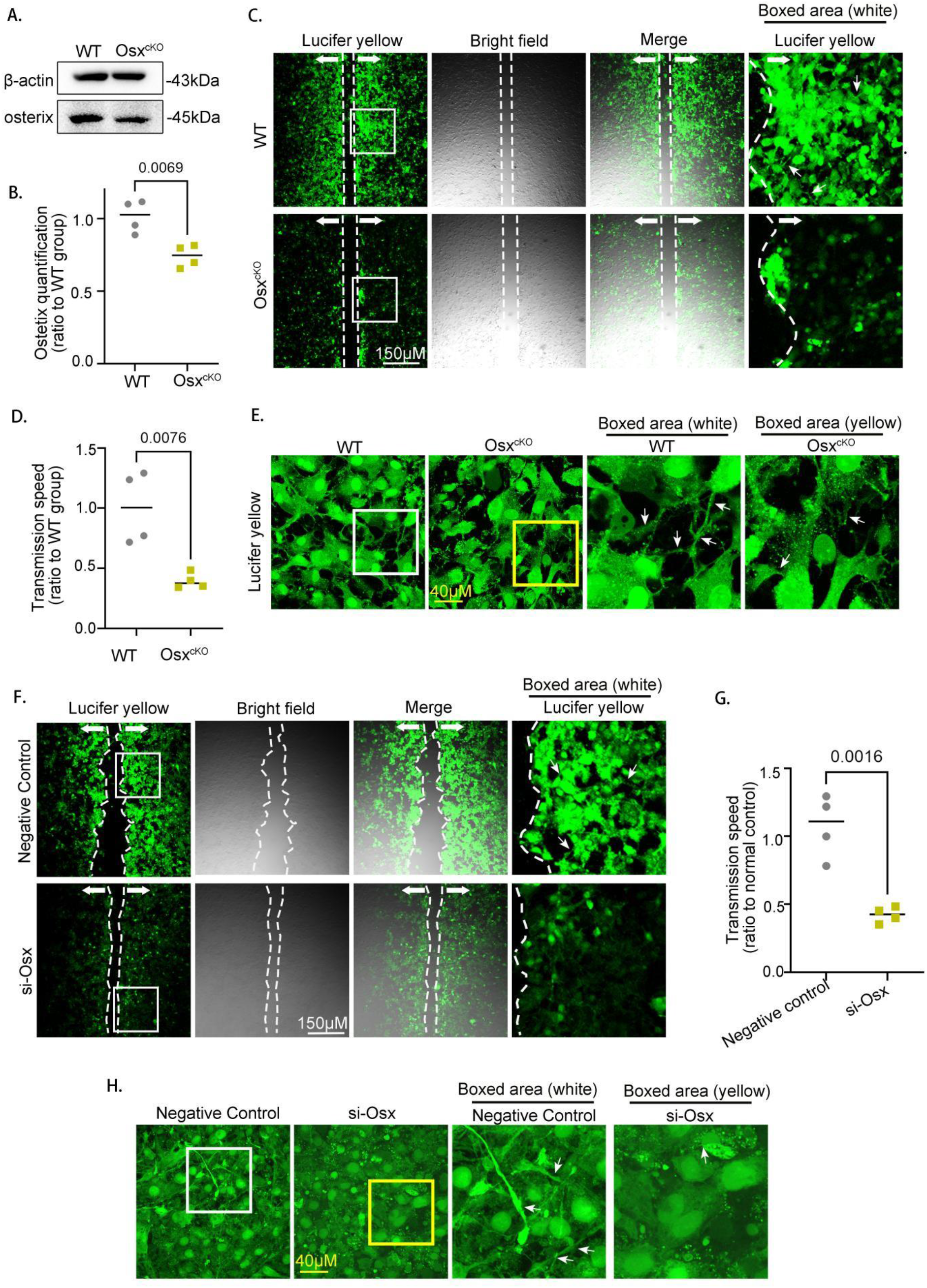
Osx deficiency compromised the transcellular fluid flow among osteocytes. (A and B) Western blots confirmed a significant reduction in Osx expression in Osx^cKO^ mice. n = 3. *p < 0.05. (C and D) Primary pre-osteocytes, differentiated from osteoblasts using osteogenic induction medium, were assessed for intercellular signaling communication using a Lucifer Yellow Dye Transfer Assay, which involved loading the cells with 1 mg/ml Lucifer Yellow dye for 10 minutes. A significant reduction in transcellular fluid flow to approximately 40% of control levels was observed in the absence of Osx. n = 3. Scale bar, 150 μM. *p < 0.05. (E) The effect shown in (C&D) was further visualized under a 60x fluorescence microscope, providing a clearer and more detailed view of the diminished gap junction channels. White arrows highlighted the reduced intercellular communication. Scale bar, 40 μM. (F and G) Mlo-y4 osteocytes were evaluated in a manner similar to (C&D), comparing the si-Osx group with the control group. n = 3. *p < 0.05. Scale bar, 150 μM. (H) The diminished gap junction in the si-Osx group was further visualized under a 60x fluorescence microscope, offering a detailed depiction of the reduced intercellular communication. White arrows indicated the areas where gap junction channels are notably decreased. Scale bar, 40 μM.

### Osx deficiency impaired cortical bone structure and decreased osteocyte dendrites both in vivo and in vitro

The immunohistochemical staining revealed that Osx was significantly knocked out in osteocytes in vivo, with a nearly 75% reduction in its positive expression as determined by statistical analysis (Fig. 2A&B). Given the strong correlation between osteocyte health and cortical bone integrity, we examined the phenotype of mouse cortical bone with a conditional knockout of Osx. We observed a marked reduction in cortical bone thickness in these mice (Fig. 2C). Considering the established link between cortical bone thickness and osteocyte dendritic function, we further investigated the role of Osx in osteocyte tibial processes. Scanning Electron Microscopy (SEM) analysis of skull and femoral sections in Osx^cKO^ mice revealed a collapse in the osteocyte dendritic network in the tibia, accompanied by diminished intercellular connections (Fig. 2D). And statistical analysis confirmed a significant decrease in the number of osteocyte dendrites (Fig. 2E). Similar findings were also observed in the skull bone, where the osteocytes’ capacity to form intercellular connections were compromised (Fig. 2F), along with a statistically significant decrease in the number of dendrites (Fig. 2G). To substantiate these results, we conducted in vitro experiments to examine the effects of Osx knockdown and overexpression on osteocytic dendrites. SEM scanning demonstrated a clear reduction in dendritic connections within osteocytes following Osx knockdown with siRNA (Fig. 2H&I). In contrast, overexpression of Osx via lentiviral vectors in osteocytes resulted in an increased number of dendritic connections (Fig. 2J&K). In summary, these comprehensive findings underscored the pivotal role of Osx in maintaining the structural integrity of the osteocyte dendritic network and its intercellular connections, which were essential for cortical bone health.

### Osx promoted the integrinαvβ1 expression and the transcellular fluid flow of osteocytes

The osteocyte dendritic network is pivotal for intercellular communication. To elucidate the impact of Osx deficiency on dendritic function, we employed the Lucifer Yellow Dye Transfer Assay (LYDTA), a method that assesses gap junction intercellular communication in osteocytes. Western blots demonstrated that the Osx expression was reduced to ∼60% compared the Osx^cKO^ group with the WT group (Fig. 3A&B). In primary pre-osteocytes, which differentiate from osteoblasts, we observed a significant reduction in transcellular fluid flow to approximately 40% of control levels in the absence of Osx (Fig. 3C&D). The effect was further visualized under a 60x fluorescence microscope, providing a clearer depiction of the phenomenon (Fig. 3E). Similar results were obtained in Mlo-y4 osteocytes following siRNA-mediated Osx knockdown (Fig. 3F- H). Collectively, these findings indicated that Osx deficiency compromises the gap junction intercellular communication of osteocytes.

Integrins, as cell adhesion molecules on the surface of vertebrate cells, are essential for intercellular connections and the anabolic response of bone tissue to mechanical stress. We hypothesized a potential relationship between dendritic changes and the integrin complex in osteocytes. Through mRNA-Seq analysis, we identified a increase in the expression of integrinαv and integrinβ1 in conjunction with Osx overexpression (Fig. 4A). This prompted us to investigate the integrinαvβ1 complex in osteocytes further. Western blots demonstrated a reduction in integrinαvβ1 protein expression in the si-Osx group compared to the control group (Fig. 4B&C). Conversely, the expression of integrinαvβ1 was significantly enhanced upon transfection of osteocytes with lenti-Osx (Fig. 4D&E). Immunofluorescence staining of osteocytes confirmed these results, showing a notable reduction of integrinαvβ1 expression in the si-Osx group and an increase in the lenti-Osx group (Figure 4F&G). In summary, Osx appeared to enhance the expression of integrinαvβ1 in osteocytes, which may underlie the observed effects on transcellular fluid flow.

**Figure 4.**
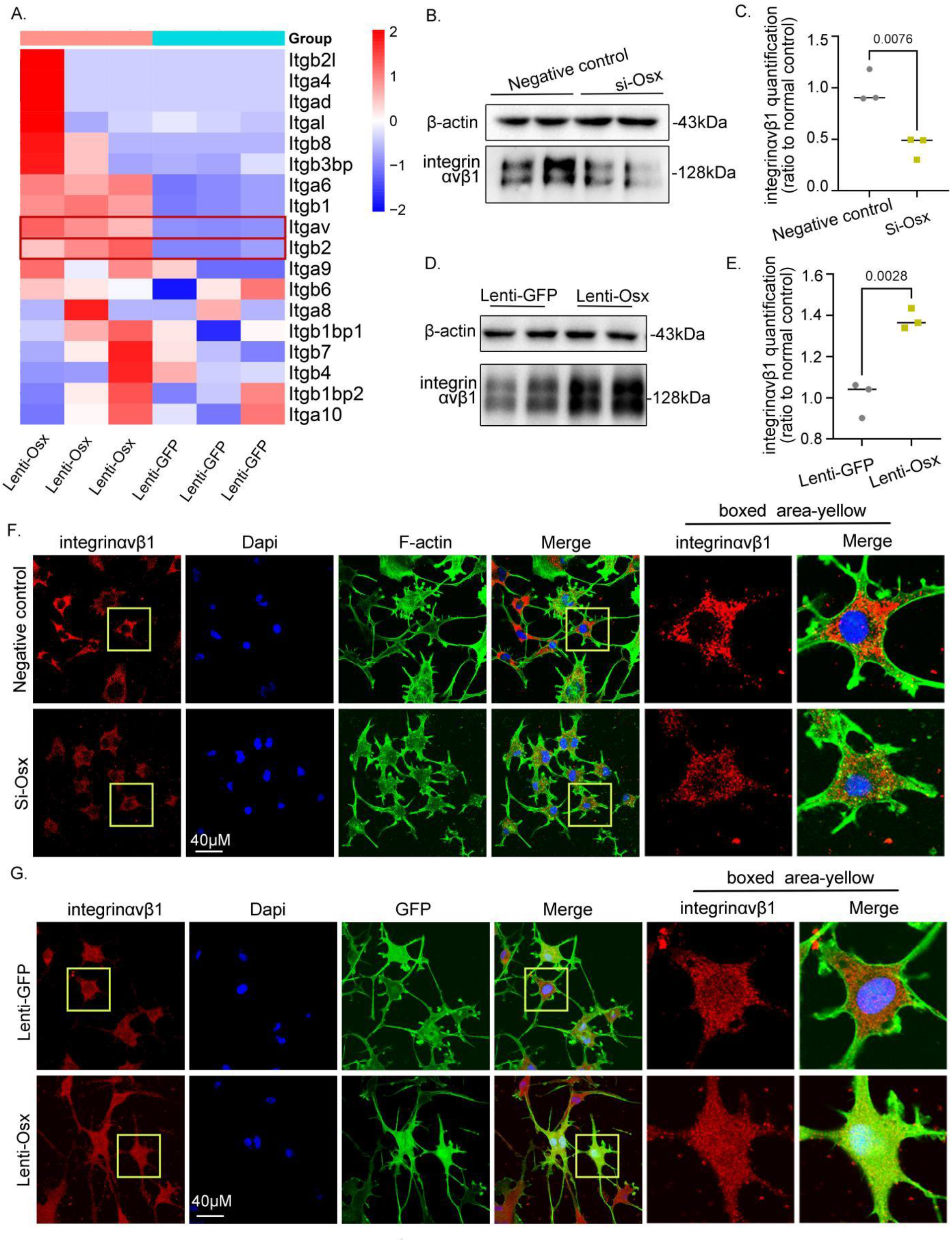
Osx deficiency reduced integrinαvβ1 expression of osteocytes. (A) A heatmap provided a comparative analysis of mRNA expression levels of the integrin family in osteocytes between the lenti-Osx group and the control group. (B and C) Western blots were used to assess the expression of integrinαvβ1 in Mlo-y4 osteocytes with si-Osx. The statistical analysis was normalized to the control group. n = 3. *p < 0.05. (D and E) Western blots were conducted to evaluate the expression of integrinαvβ1 in Mlo-y4 osteocytes with lenti-Osx. The statistical analysis was normalized to the control group. n = 3. *p < 0.05. (F) Immunofluorescence staining illustrated a reduction in integrinαvβ1 expression at both the osteocyte body and its intercellular dendrites with Osx deficiency. The cytoskeleton was stained green by F-actin, integrinαvβ1 was stained red, and nuclei were stained blue. n = 3. Scale bar, 40 μM. (G) Immunofluorescence staining conversely showed an increase in integrinαvβ1 expression at the osteocyte body and its intercellular dendrites with the introduction of Lenti-Osx. The cytoskeleton, integrinαvβ1, and nuclei were stained as described in (F). n = 3. Scale bar, 40 μM.

### Cx43 was the transcriptional target of Osx in maintaining osteocytic dendrites

Osx is usually known as a pivotal transcription factor for bone formation, and has been verified above to implicating in the collapse of the intercellular dendritic network and the diminished intercellular communication among osteocytes. Thus, to elucidate the downstream targets of Osx, we conducted a comprehensive analysis using CHIP-seq, focusing on the overexpression of Osx in osteocytes via lentiviral transfection. Our findings revealed significant differences in expression peaks between the lentivirus-Osx (lenti-Osx) group and the control group, which received the lentivirus-GFP vector (lenti-GFP) (Fig. 5A&B). Gene Ontology (GO) analysis showed that Osx predominantly enhanced various signaling pathways in osteocytes, such as "synapse," "dendrite development," "neuron projection," "dendrite," "cell projection," and "cell junction" (Fig. 5B). Furthermore, GO analysis of mRNA-seq data suggested that Osx influences cell synapses and cytoskeleton functions, including "microtubule motor activity," "structural constituent of cytoskeleton," "microtubule end," "cell synapse," and "integrin complex" (Fig. 5C). Our investigation was particularly focused on the GJA family of genes, known for their critical role in intercellular communication. Our data indicated that overexpression of Osx increased the expression of GJA1(Cx43), whereas those of other GJA family members did not exhibit obvious changes (Fig. 5D). To further investigate the regulatory mechanism, we identified two distinct peaks in the Cx43 promoter between the lentivirus-GFP and lentivirus-Osx groups through CHIP-seq analysis (Fig. 5E), suggesting a direct binding of Osx to the promoter of the GJA1 gene, thus regulating its transcription.

**Figure 5.**
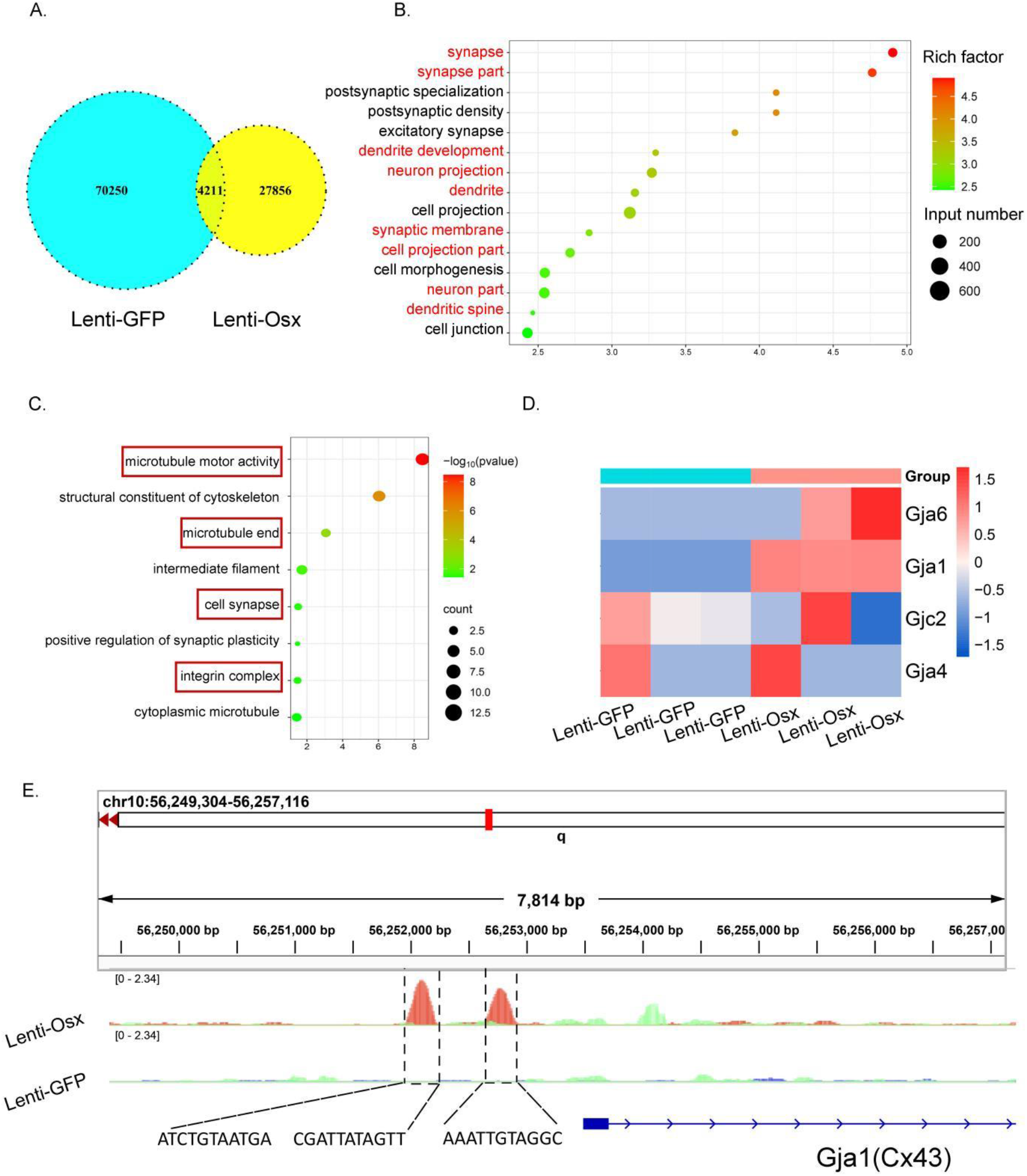
Cx43 was the transcriptional target of Osx in osteocytes. (A) The plot displayed the differences in expression peaks between the lenti-Osx group and the control group. (B) Gene Ontology (GO) analysis was conducted on the differential expression peaks identified after performing CHIP-seq experiments with Osx. The analysis suggested that Osx played a regulatory role in multiple signaling pathways within osteocytes. (C) GO analysis of mRNA-seq data indicated that the functions of Osx were predominantly associated with dendrite-related processes in osteocytes. (D) A heatmap illustrated the changes in expression levels of the gap junction protein family (GJA) in osteocytes with Lenti-Osx as determined by mRNA-seq analysis. (E) CHIP-seq analysis identified two distinct peaks in the Cx43 promoter region binding with Osx, highlighting the direct transcriptional regulation of Cx43 by Osx.

Then we further to confirmed Cx43 as a downstream target of Osx both in vivo and in vitro. In vivo immunohistochemical staining showed a significant reduction in Cx43 expression within the osteocytes of bone lacunae of Osx^cKO^ mice compared to the control (Fig. 6A & B), and which was also confirmed using vivo immunofluorescence staining (fig. S1A). Using primary pre-osteocytes derived from Osx^cKO^ mice, demonstrated a reduction in Cx43 expression, both at the protein and mRNA levels by WB and qPCR analysis (Fig. 6C-E). Immunofluorescence analysis for cell culture also indicated diminished Cx43 expression in primary pre-osteocytes of the Osx^cKO^ group at gap junctions (Fig. 6F). These findings were further corroborated by overexpression of Osx in osteocytes via lentiviral transfection led to an increasement of Cx43 expression at both protein and mRNA levels, using Western blots and quantitative real-time PCR analysis (Fig. 6G-J). Confocal immunofluorescence analysis also revealed that osteocytes transfected with lenti-Osx displayed robust Cx43 expression at sites where dendritic processes connect, forming an extensive intercellular dendritic network, distinct from the control group’s patchy linear distribution (Fig. 6K). In addition, Cx43 expression in Mlo-y4 osteocytes treated with Si-Osx was reduced by half at both protein and mRNA levels compared to the control group, as shown by WB and qPCR analysis (fig. S1B-D), with further confirmation by immunofluorescence staining (fig. S1E). In contrast to the control group’s linear red fluorescence marking Cx43 expression along dendritic processes, the Si-Osx group exhibited a scattered, dot-like distribution of Cx43 between the dendritic processes of osteocytes (fig. S1E). Collectively, this comprehensive results elucidated the direct regulatory role of Osx in the transcription of Cx43 and its subsequent impact on intercellular communication in osteocytes.

**Figure 6.**
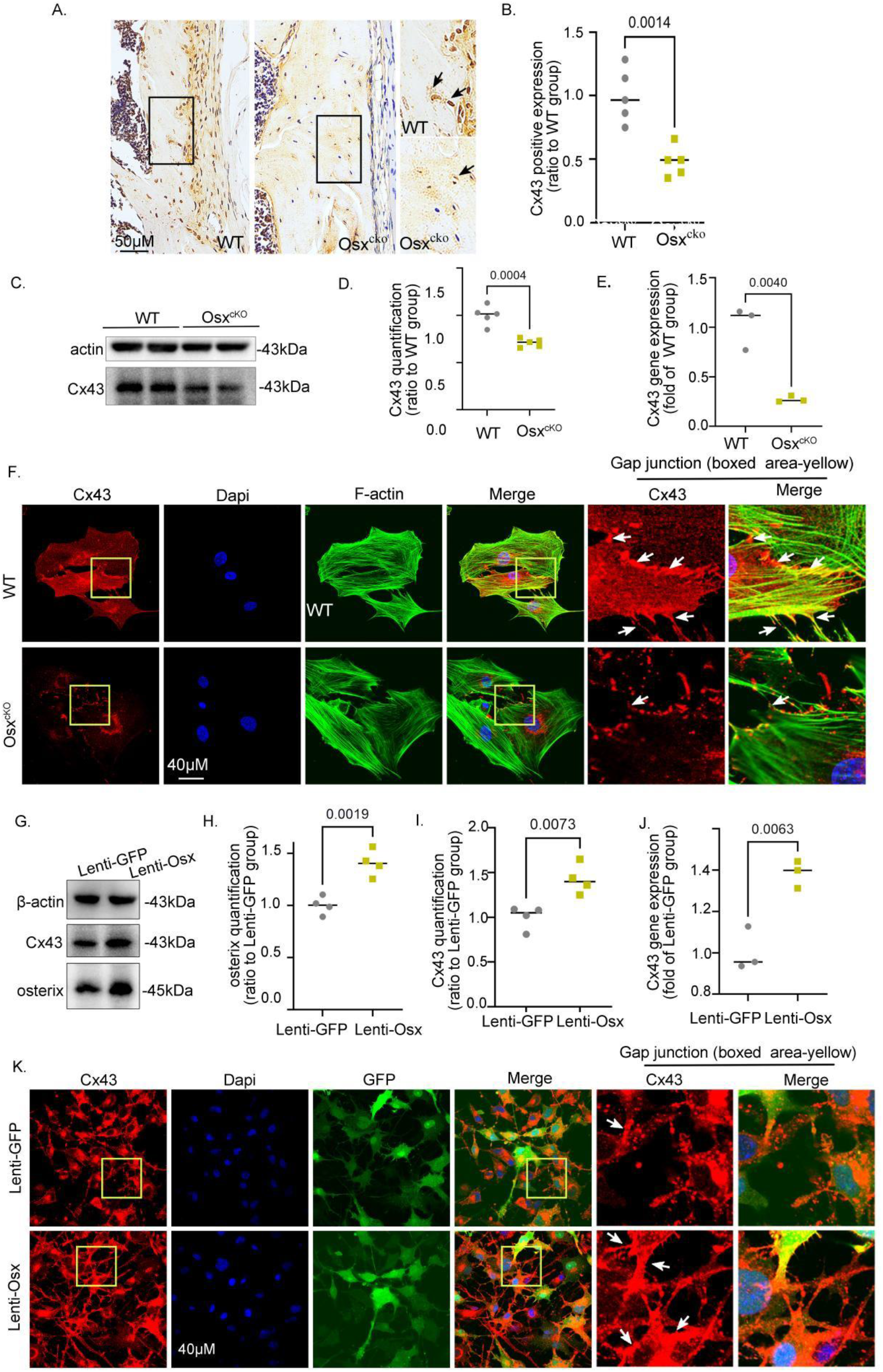
Osx promoted Cx43 expresion of osteocytes both in vitro and in vivo. (A and B) In vivo immunohistochemical staining revealed a diminished Cx43 expression in the cortical bone of Osx^cKO^ mice as opposed to the WT mice. Statistical analysis was performed using Image J software with normalization to the control group. Scale bar, 50 μM. *, p < 0.05. (C and D) Western blots showed a decrease in Cx43 expression of Osx^cKO^ pre- osteocytes. n = 3. Statistical analysis was based on the control group after normalization. *p < 0.05. (E) qPCR showed the mRNA levels of Cx43 in Osx^cKO^ pre-osteocytes. n = 3. *p < 0.05. (F) Immunofluorescence staining depicted a diminished expression of Cx43 in Osx^cKO^ pre-osteocytes relative to the WT group. The cytoskeleton (F-actin) was visualized in green, Cx43 in red, and nuclei in blue. n = 3. Scale bar, 40 μM. (G, H and I) Western blots showed Cx43 expression level in Mlo-y4 osteocytes with lenti-Osx. Statistical comparisons were made relative to the control group after normalization. n=3. *p < 0.05. (J) qPCR corroborated the increase of Cx43 mRNA in Mlo-y4 osteocytes with lenti-Osx. n=3. *p < 0.05. (K) Immunofluorescence staining demonstrated an elevated Cx43 expression at the gap junction of osteocytes in the Lenti-Osx group (L) compared to the WT group. The images showed GFP in green, Cx43 in red, and nuclei in blue. n=3. Scale bar, 40 μM.

### Restoration of Cx43 rescued intercellular communication among Osx-deficient osteocytes both in vivo and in vitro

To solidly elucidate the role of the Osx-Cx43 axis for osteocytic communication, we conducted rescue experiments both in vivo and in vitro. We aimed to restore Cx43 signaling by using all-trans-retinoic acid (ATRA), a known agonist of Cx43. Western blots revealed that treatment with ATRA significantly increased Cx43 expression (Fig. 7A). When Mlo-y4 osteocytes, previously transfected with si-Osx, were treated with ATRA, we observed a significant recovery in both the reduced cellular migration and the impaired trans-cellular fluid flow, as indicated by the Lucifer yellow dye (Fig. 7B&C). In vivo, ATRA was administered to the cranial bone of mice, and we found that the dendritic connections among osteocytes in the Osx^cKO^ + ATRA group were notably enhanced compared to the Osx^cKO^ group (Fig. 7D&E). In addition, both Western blots and fluorescence staining demonstrated that the expression of integrinαvβ1, which was inhibited by si-Osx, was also reversed by ATRA treatment (Fig. 7F&G). These data collectively demonstrated that restoring Cx43 signaling through ATRA effectively reversed the impaired communication in osteocytes caused by Osx deficiency, both in vitro and in vivo.

**Figure 7.**
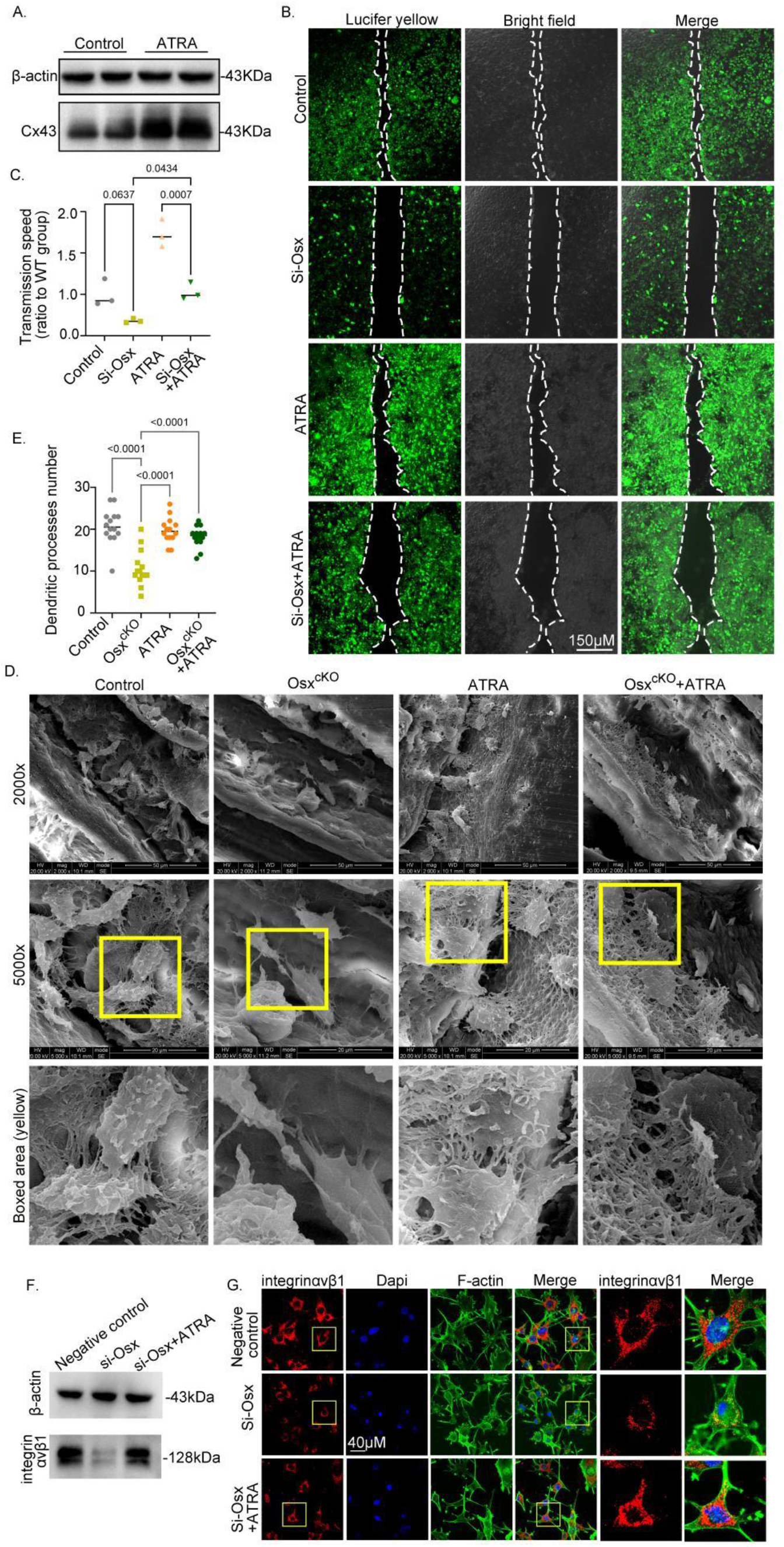
Restoration of Cx43 rescued intercellular communication among Osx- deficient osteocytes both in vivo and in vitro. (A) Western blots was employed to assess the impact of all-trans-retinoic acid (ATRA), a Cx43 signaling agonist, on Cx43 expression in Mlo-y4 osteocytes. n = 3. (B and C) The Lucifer Yellow Dye Transfer Assay was utilized to evaluate the gap junction intercellular communication among osteocytes. ATRA intervention significantly restored the impaired communication observed in Osx-deficient osteocytes. n = 3. Scale bar, 150 μM. (D and E) Scanning electron microscopy (SEM) was conducted to visualize the osteocytic dendrites in the cranial bone. The results demonstrated that ATRA intervention effectively reversed the reduction in dendritic density of osteocytes observed in Osx^cKO^ mice. Dendrite counts were performed per osteocyte, with 16 osteocytes analyzed per group. Scale bar, 50 μM & 20 μM. *p < 0.05. (F) Western blots revealed a decrease in integrinαvβ1 expression in Mlo-y4 osteocytes with si-Osx. By contrast, ATRA treatment rescued the reduced integrinαvβ1 expression. (G) Immunofluorescent staining corroborated the Western blots findings, showing that ATRA intervention restored integrinαvβ1 expression in osteocytes with si-Osx. The cytoskeleton (F-actin) was visualized in green, integrinαvβ1 in red, and nuclei in blue. Scale bar, 40 μM.

**Figure 8.**
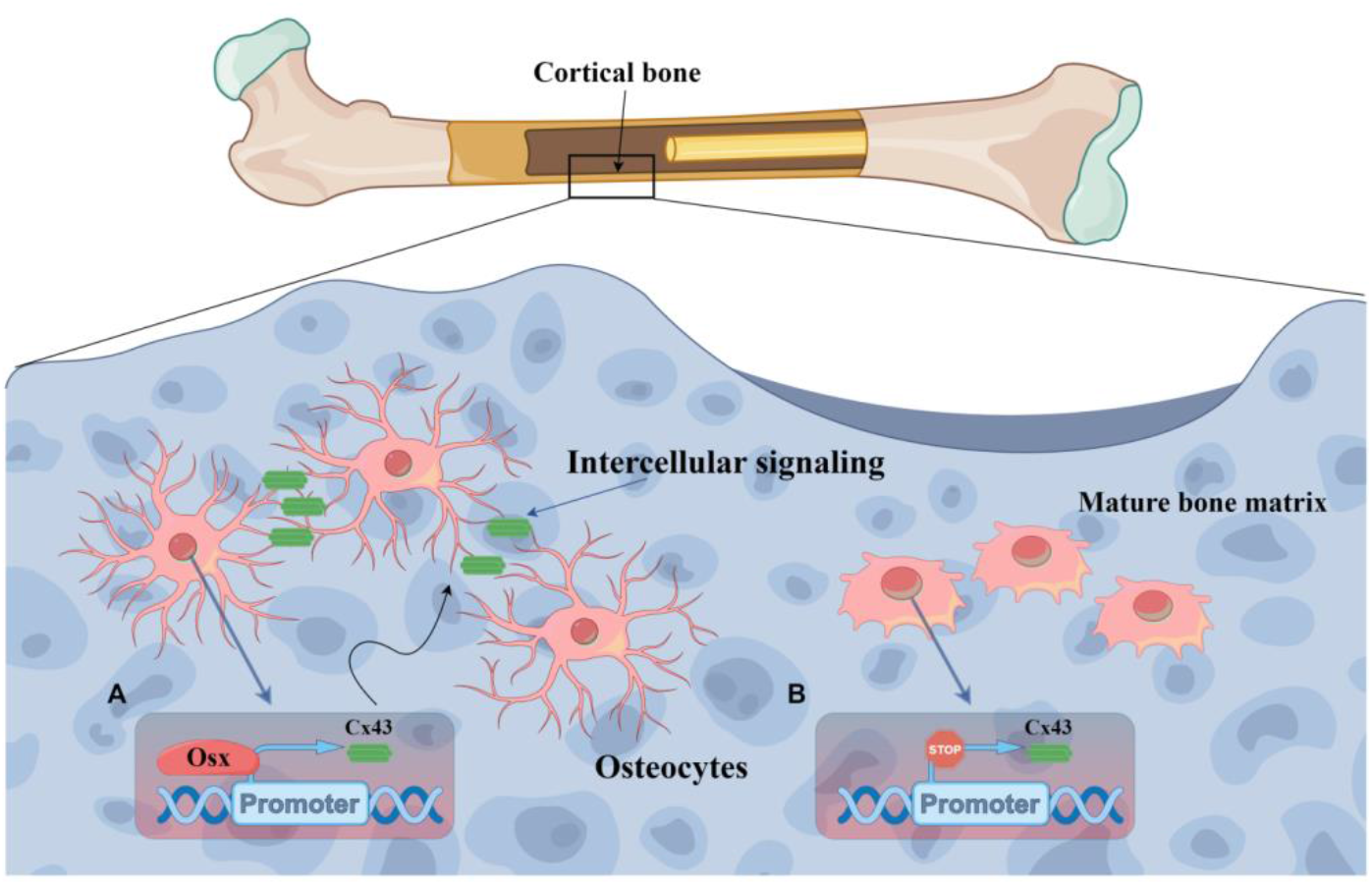
Schematic model illustrated the Osx-Cx43 axis of osteocytes and its role in intercellular signaling communication. The dendrites of osteocytes were essential for maintaining gap junction intercellular communication, given that osteocytes were non-migratory, as embedded within the bone mineralized matrix. **(A)** In physiological state, Osx directly promoted the transcription of Cx43, the key component of gap junction channel, which was a critical regulatory mechanism that preserved the integrity and function of osteocytic dendrites, thus governing the transmission of signals among osteocytes within the mature bone tissue. **(B)** In pathological state, Osx deficiency inhibited the Cx43 expression, then restrained the dendritic network among osteocytes, thereby impairing osteocyte in response to environmental stimuli and signal transduction, such as mechanical forces or bone fracture.

## Discussion

Bone remodeling in response to internal and external environmental stimuli necessitates the cellular signal communication of osteocytes, osteoblastes, and osteoclasts, ensuring the mechanical competence and structural integrity. Although the roles of osteoblasts and osteoclasts in bone remodeling are extensively understood, osteocytes in controlling the bone remodeling response need to be studied in depth. Adult osteocyte dysfunction was found to cause increased cortical bone porosity, deterioration of the bone microstructure, medullary cavity fat hyperplasia, loss of trabecular bone, and microfractures(Plotkin & Bellido, 2016). Osteocytes possess gap junctions, which are narrow channels extending to neighboring cells, allowing communication through the transfer of small molecules and ions. Connexins are key molecules for cell-cell connections and intercellular communication among osteocytes(Qin et al., 2020). When the dendrites of osteocytes receive mechanical stimuli, integrins and connexins in osteocytes get activated, thereby promoting the further transmission of mechanical signals(Riquelme et al., 2021). Therefore, it is important to further explore the specific mechanisms of intercellular communication among osteocytes for understanding bone remodeling and the related diseases. This study uncovered a distinct regulatory mechanism of osteocytic communication that was linked to gap junctions.

Traditionally perceived, Osx is known as a specific regulator of osteoblast differentiation during new bone formation, essential for maintaining bone homeostasis in mammals. Osteogenesis imperfecta type 12 (OI12) is caused by monoallelic or biallelic mutations in the Osx gene. It is characterized by recurrent fractures, skeletal deformities, and is closely associated with bone fragility(Gauthier et al., 2023). Postnatal deactivation of Osx in mice leads to multiple skeletal abnormalities, including the absence of new bone formation, failure to resorb mineralized cartilage, and defects in bone cell maturation and function(Sinha & Zhou, 2013; Zhou et al., 2010). While previous studies focus predominantly on the role of Osx in osteoblast differentiation and bone formation, the role of Osx in osteocytes is not yet fully comprehended. This study demonstrated a distinct function of Osx. Osx deficiency in osteocytes leaded to a reduction in the formation of intercellular dendritic processes and a collapse of the dendritic network structure. This findings underscored the uniqueness of this intercellular signaling network to osteocytes, and potentially illuminating the microscopic processes involved in bone dysplasia. The Lucifer Yellow Dye Transfer Assay confirmed that Osx is directly linked to intercellular communication in osteocytes. The communication capability of osteocytes diminished and the adhesion-related molecule integrin reduced with Osx deficiency. Subsequently, we identified target genes of Osx that govern the dendritic network by mRNA-seq. Gap junctions, primarily responsible for intercellular communication, involve the connexin family, which forms hexameric hemichannels on the cell membrane(Ma et al., 2019). Connexins are the most abundant gap junction proteins expressed in osteocytes(Hoppock et al., 2023). A hemichannel composed of Cx43 is present on the cell processes of osteocytes. This hemichannel opens in response to mechanical stimulation, allowing the release of small molecular metabolic substances, such as prostaglandin E2 (PGE2)(Hoppock et al., 2023). Hemichannels from adjacent osteocytes combine to form gap junction channels, allowing for the exchange of siganling like ions or secondary messenger molecules between two neighboring osteocytes(Ma et al., 2019). Our systematic mRNA sequencing analysis of connexin family expression revealed that Cx43 (GJA1) exhibits the most significant expression difference within the GJA family. Western blots, immunofluorescence, and immunohistochemical staining together indicated that Osx deficiency directly affected the expression of Cx43, which was essential for bone development and maintenance, particularly in cortical bone(Moorer & Stains, 2017).

Previous studies have indicated that Cx43 is crucial for cortical bone thickness, with its targeted deletion in osteocytes leading to increased apoptosis of osteocytes and enhanced recruitment of osteoclasts(Epifantseva et al., 2020; Lloyd et al., 2012; Moorer & Stains, 2017). Mutations of Cx43 are associated with various disorders, including Oculo-dento-digital dysplasia (ODDD), hypoplastic left heart syndrome, craniofacial malformation syndrome, and atrioventricular septal defect(Delmar et al., 2018). The absence of Cx43 in osteocytes can delay healing or lead to non-union of fractures,(Chen et al., 2019) and result in structural changes, including thinning of the bone cortex and signs of bone aging, which align with the functions of Osx(Lloyd et al., 2012; Moriishi et al., 2022). Given the potential overlapping roles of Cx43 and the phenotyes in this study on Osx, we hypothesized that they shared functional relationship in osteocytes. To test this, we conducted in vivo and in vitro experiments that demonstrated a decrease in Cx43 expression in the bone cortex following Osx deficiency. Our study conclusively showed that Cx43 acted downstream of Osx, regulating the intercellular dendritic network and cell-cell communication. The relationship between Osx and Cx43 was further explored through CHIP-seq analysis, which showed Osx was enriched in the promoter region of Cx43. It’s also reported that the suppression of Cx43 expression by shRNA against Osx in HEK293 and C2C12 cells, and its enhancement through Osx overexpression are observed, which is align with our findings.(Han et al., 2016) Moreover, ATRA, acting as an agonist of Cx43, facilitates intercellular communication among osteocytes. Our previous work has convincingly demonstrated ATRA’s role in enhancing intercellular communication among periodontal ligament cells(Wu et al., 2022). In vivo, after administering ATRA into the skull, we harvested cranial bone tissue and found that the dendritic network of osteocyte, which was diminished in Osx deficiency mice, was reversed by ATRA treatment. These findings collectively suggested that Cx43 acts as a direct target of Osx, and involved in the regulatory of osteocyte dendritic network.

It’s worth noting that in the experiments using primary osteocytes for in vitro cell culture, we couldn’t entirely eliminate the presence of osteoblasts as a potential distraction, although these primary cells were cultured by osteogenic induction into osteocytes. Consequently, the resulting data may reflect a mixture of osteoblasts, pre-osteocytes, and osteocytes. The regulatory role of Osx in osteoblast differentiation appears to indirectly promote its function in osteocytic communication, though this requires further clarification. Nevertheless, these findings consistently pointed to Osx playing a distinct role in enhancing intercellular signaling among osteocytes by preserving their dendrites. Furthermore, we demonstrated that Osx directly promoted Cx43 signaling at the transcriptional level, elucidating a new mechanism different from osteoblast differentiation. These insights were valuable for developing strategies aimed at addressing conditions such as fracture nonunion or disuse osteoporosis in bone diseases.

## Methods

### Mice

All animal procedures were approved by the Animal Care and Use Committee of Sichuan University. Mice were meticulously housed at room temperature (23°) in the animal room at West China Hospital of Sichuan University. Mice had free access to food and water at all times and were maintained under a 12-hour alternating light-dark cycle and the humidity levels meticulously maintained between 50% and 60% in specific-pathogen free conditions (SPF), ensuring optimal living conditions. The Col1α1-CreER mice and the Osx^flox/flox^ mice were bought from GemPharmatech (Nanjing, China), and were crossed with each other to generate Osx^cKO^ mice (Osx^flox/flox^;Col1α1-CreER), and the wild type littermates (Osx^flox/flox^) as controls (WT). Both Osx^cKO^ mice and their littermate controls received an intraperitoneal injection of tamoxifen at a dosage rate of 0.1 mg per gram of body weight, and corn oil was used as the solvent. This treatment was administered for six consecutive days. In the experimental setup aimed at assessing the influence of the Cx43 agonist ATRA, the mice were treated with skull injections of ATRA at the age of four weeks, with the injections continuing for a period of two weeks. The collection of skull samples was conducted at the age of eight weeks for subsequent analysis.

### Preparation of tissues and cells

Mouse osteocyte-like Mlo-y4 cells were sourced from the American Type Culture Collection (Manassas, VA), in accordance with established protocols.(Liu et al., 2019) These cells were cultured following 4 to 6 passages in Alpha’s Modified Eagle Medium (αMEM), obtained from Gibco/Life Technologies (Carlsbad, CA, USA) , and mixed with 10% fetal bovine serum (FBS). The Mlo-y4 cells were plated at a density of 3×10^5^ cells per well in 6-well plates and incubated at 37°C in a controlled atmosphere containing 5% CO_2_. The extraction of primary pre-osteocytes/osteocytes was performed as detailed in "*Bone Research Protocols*" by Gooi JH *et al*., published in Methods Mol Biol, 2019. The process began with the aseptic dissection of femurs and tibias from 6-week-old mice. All extraneous muscle, tendons, and periosteum were carefully removed in alpha-Minimum Essential Medium (α-MEM), supplemented with 10% penicillin and streptomycin, using product number SH30265.01 from HyClone (Logan, USA). The epiphyses were then severed, and bone marrow was expelled using a syringe. The bones were cut into 1-2 mm segments and subjected to 2-3 wash cycles in α-MEM containing 2% penicillin and streptomycin. The bone fragments were incubated for 30 minutes at 37°C in α-MEM enriched with 10 active units/ml of collagenase II, sourced from Gibco (Grand Island, USA). This was followed by triple washing with phosphate-buffered saline (PBS). Subsequently, the fragments were treated with a 5 mM solution of ethylenediaminetetraacetic acid (EDTA) from Macklin (Shanghai, China), for another 30 minutes at 37°C, followed by washing with PBS. The bone fragments underwent two alternating incubation cycles with collagenase II and EDTA. Finally, the residual fragments were mechanically disaggregated in α-MEM using a tissue homogenizer. The resulting bone particulate suspension was directly seeded onto plates pre-coated with 4% type I collagen from Corning (New York, USA). The cultures were maintained for seven days in α-MEM supplemented with 5% FBS from Summit Biotechnology (Fort Collins, CO, USA), and 5% calf serum from HyClone Laboratories (Logan, UT, USA), to facilitate the migration of osteocytes from the bone fragments.

### Real time - Quantitative PCR (RT-qPCR)

Total mRNA was meticulously isolated from osteocytes using the RNeasy RNA Purification Kit from Qiagen, adhering to the manufacturer’s instructions. Thereafter, 1 µg of RNA was reverse transcribed into complementary DNA (cDNA) using the iScript cDNA Reverse Transcription Kit (R423-01, Va, China), following the manufacturer’s guidelines. RT-qPCR was performed on the iCycler platform from Bio-Rad in the USA, with each reaction comprising 1 μL of template cDNA within a 25 µL total volume. The primer sequences, detailed in Table 1, were designed with glyceraldehyde-3-phosphate dehydrogenase (GAPDH) serving as the reference control. The relative mRNA levels were ascertained using the comparative threshold cycle method, a standard approach in quantifying gene expression levels.

### Osx small interfering RNA (siRNA) transfection

Osteocytes were plated into 3.5mm cell culture dishes and allowed to settle for a period of 12 hours. Following this incubation, the siRNA transfection complex was prepared using RNAiMAX (Invitrogen, California, USA), and added to each well at the volume specified by the manufacturer’s instructions. The culture plate was gently agitated to ensure complete integration of the transfection complex with the cells. The specific sequences of the siRNA employed were as follows: for the sense strand, 5’- *CAAGAUGUCUAUAAGCCCATT*-3’; and for the antisense strand, 5’- *UGGGCUUAUAGACAUCUUGTT*-3’. A non-targeting RNA sequence was utilized as a control for the scrambled group to ensure the specificity of the siRNA effect. At 48 hours post-transfection, both RT-qPCR and Western blots were conducted to evaluate the mRNA and protein levels of Osx and Cx43, respectively. These analysis provide a comprehensive assessment of the efficiency of siRNA-mediated gene knockdown and its subsequent impact on the expression of target genes.

### Lentiviral transfection of Osx

The mouse Osx overexpression lentiviral vector, designated as HBLV-m-Osx-3xflag- ZsGreen-PURO, was expertly packaged and constructed by HanBio Technology (Shanghai, China) and introduced into Mlo-y4 osteocytes. The integrity of this construct was meticulously confirmed through PCR analysis. A control vector, HBLV-ZsGreen-PURO, was similarly packaged for comparative studies. For the transfection process, Mlo-y4 cells were prepared using a Lipofectamine Transfection Kit, applied at a density of 1 × 10^5^ cells/mL and a viral titer of 1 × 10^8^ TU/mL, strictly adhering to the manufacturer’s instructions.

### Western blots

Prior to analysis, osteocytes were gently washed twice with precooled Phosphate-Buffered Saline (PBS) and harvested using a cell scraper. The cells were then lysed on ice for 10 minutes with RIPA Cell Lysis Buffer (Beyotime, P0013B, China), followed by high-temperature denaturation to prepare for immunoblotting. Proteins were resolved by Sodium Dodecyl Sulfate-Polyacrylamide Gel Electrophoresis (SDS-PAGE) and transferred onto Polyvinylidene Difluoride (PVDF) membranes. The membranes were initially blocked with 5% skim milk powder, followed by an overnight incubation at 4°C with primary antibodies specific for Osx and Cx43 both anti-rabbit (Abcam, Cambridge, UK, catalog numbers ab209484 and ab78055, respectively). On the subsequent day, the membranes were incubated with a secondary antibody (GB23302, Wuhan Servicebio Co., Ltd., China) for 2 hours at room temperature. Detection of the proteins of interest, Cx43 and Osx, was achieved using an Enhanced Chemiluminescence (ECL) detection kit. The expression levels of these proteins were quantified using Image J software (NIH, Bethesda, MD) and normalized against β-actin to ensure the accuracy and reliability of the results.

### Scanning electron microscopy (SEM)

To discern alterations in the cortical and cranial dendritic networks of mice, specimens were harvested at precise time intervals. The collected samples were initially fixed in a 70% ethanol solution for a duration of 48 hours, followed by a process of gradient dehydration. Subsequently, the samples were embedded in polymethyl methacrylate (PMMA), a common resin used for its excellent preservation properties. The bone tissue sections were meticulously polished using an automatic grinding system (Exakt, Germany). The system operated at varying rotational speeds to achieve a smooth surface finish. The sections were then subjected to a brief immersion in a 19% phosphoric acid solution for exactly 5 minutes, which was immediately followed by a thorough rinse in deionized water to neutralize the acid. Further treatment involved a 20-minute immersion in a 5% sodium hypochlorite solution, a step critical for enhancing the contrast of the bone’s surface features. After air-drying, the bone tissue surface was meticulously coated with a thin layer of gold alloy to optimize conductivity and reduce charging during SEM examination. The specimens were then examined using SEM, a high-resolution imaging technique that provided detailed visualization of the bone’s microstructure and dendritic networks, revealing the intricate morphological changes.

### Scrape loading and dye transfer

The Lucifer Yellow Scratch Stain experiment stands as a widely recognized method for the detection of intercellular communication. This technique involves creating a scratch with a blade, facilitating the transfer of a fluorescent yellow dye to distant cells along the ones bordering the scratch. To evaluate the impact of Osx expression on inter- osteocyte communication, cells were engaged in fluorescent dye transfer experiments. Upon reaching full confluence, the cell monolayer was carefully incised with a blade. The monolayer was then rinsed with Calcium and Magnesium-free Phosphate-Buffered Saline (CAMG-PBS) to prepare the cells for dye uptake. Immediately post-incision, the monolayer was stained with a fluorescent dye solution at a concentration of 1 mg/ml. Following a brief incubation period of 2 minutes at room temperature, the cells were gently washed to eliminate any surplus stain. The subsequent transfer of the fluorescent dye was meticulously monitored using a confocal microscope over a duration of 7 minutes. This observation period allowed for the assessment of the communication pathways between osteocytes, providing insights into the role of Osx in the connectivity and communication efficiency within the cellular network.

### Immunohistochemistry and immunofluorescence

Immunohistochemistry (IHC) and immunofluorescence (IF) analysis were conducted following established protocols. After the initial hydration process, tissue sections underwent antigen retrieval in a citric acid buffer solution with a concentration of 10 mM at a pH of 6.0. For IHC staining, the sections were first incubated in 3% hydrogen peroxide in the dark for 25 minutes to quench endogenous peroxidase activity. This was followed by a blocking step with 10% rabbit serum for 30 minutes to minimize non-specific binding. The sections were then incubated with the primary antibody diluted in 1% Bovine Serum Albumin (BSA) overnight at 4°C. Subsequently, the sections were treated with a biotinylated secondary antibody, diluted 1:200, for 2 hours at room temperature. Positive immunoreactivity was visualized using the chromogenic substrate 3,3’-diaminobenzidine (DAB), while cell nuclei were counterstained with hematoxylin for enhanced contrast. The extent of positive staining in osteocytes was meticulously quantified using Image-Pro Plus 6.0 software from Media Cybernetics (Maryland, USA). For IF staining, following incubation with the primary antibody, the sections or osteocytes cultured on confocal dishes were treated with secondary antibodies conjugated to Alexa Fluor 647, a far-red fluorescent protein tag, from Life Technologies (Carlsbad, CA, USA). This incubation was performed for 1 hour at room temperature in the dark to prevent photo-bleaching. Cell nuclei were stained with 4’,6-diamidino-2- phenylindole (DAPI), a fluorescent stain specific for DNA, from Thermo Fisher Scientific (Waltham, MA, USA). The cytoskeleton was visualized by staining with phalloidin, a potent actin filament marker, at a concentration of 6 μM (Invitrogen, California, USA). The resulting fluorescent signals were captured using a high-resolution FluoView FV1000 confocal microscope from Olympus. The images were further analyzed using FV10-ASW Viewer software version 4.2 and ImageJ version 1.8, both of which are widely utilized for advanced image processing and analysis in biological research.

### RNA sequencing

Total mRNA was meticulously extracted from the samples using Trizol reagent (Invitrogen, California, USA), following the manufacturer’s recommended protocol. The quality of the extracted mRNA was rigorously assessed utilizing a Nanodrop™ OneC spectrophotometer (Thermo Fisher Scientific Inc, USA), ensuring the purity and concentration of the mRNA preparations. To further ensure the integrity of the mRNA, samples were subjected to RNase-free agarose gel electrophoresis, a standard method for visualizing mRNA integrity and size distribution. Following this, the mRNA was reverse transcribed to generate complementary DNA (cDNA), which served as the template for subsequent PCR amplification. The samples were prepared for PCR amplification on the Illumina NovaSeq 6000 platform (Contest Biotechnology, China), a high-throughput sequencing system allowing for the generation of vast amounts of sequencing data. Gene Ontology (GO) enrichment analysis was conducted to identify significantly enriched biological processes, molecular functions, and cellular components among the differentially expressed genes. This analysis was performed using the GOseq R software package in conjunction with the DAVID online tool, available at https://david.ncifcrf.gov/. Additionally, pathway analysis of the differentially expressed genes was carried out to explore the biological pathways they are involved in. This was performed using the Kyoto Encyclopedia of Genes and Genomes (KEGG) database, accessible at http://www.kegg.jp/KEGG/, and the Gene Set Enrichment Analysis (GSEA) software, version 4.0.3, to identify pathways with significant changes in gene expression patterns. These comprehensive analysis provide a robust framework for understanding the molecular mechanisms underlying the observed phenotypes and the role of specific genes in the biological context of the study.

### CHIP-seq analysis

The CHIP assay was expertly executed by SeqHealth (Wuhan, China). Osteocytes were fixed utilizing a 1% formaldehyde solution for a duration of 10 minutes at room temperature, a critical step to promote protein-DNA cross-linking. To terminate the cross-linking reaction, 125 mM glycine was introduced and the mixture was allowed to stand for 5 minutes. Following this, the cells were harvested and rapidly frozen in liquid nitrogen to preserve the chromatin structure. Micrococcal nuclease (MNase) was then employed to digest the chromatin, fragmenting the DNA into pieces that ranged from 200 to 600 base pairs (bp) in length. These chromatin fragments were subjected to an overnight incubation with an anti-Flag antibody (F1804, Sigma-Aldrich) to carry out immunoprecipitation. The cross-links of the chromatin DNA were reversed through de- crosslinking at 55°C in conjunction with Proteinase K for a period of 5 hours, which effectively prepared the DNA for downstream sequencing processes. Both the CHIP (Chromatin Immunoprecipitation) and input DNA libraries were sequenced using the high-throughput NovaSeq 6000 sequencer from Illumina. This CHIP-seq analysis provides a detailed map of protein-DNA interactions within the genome, offering valuable insights into the regulatory mechanisms that govern gene expression in osteocytes.

### Statistical analysis

The experiments were meticulously carried out in triplicate to ensure the reliability and reproducibility of the results. Quantitative data are presented as the mean ± standard deviation (SD), a standard measure of statistical dispersion that provides an indication of the variability within the dataset. These results were graphically represented using GraphPad Prism (GraphPad Prism Inc., San Diego, USA), renowned for its capabilities in data visualization and analysis. For the statistical evaluation of the data, one-way Analysis of Variance (ANOVA) was employed to determine the presence of significant differences among group means. This initial test was followed by Tukey’s Honest Significant Difference (HSD) post-hoctest, a widely accepted method for pairwise multiple comparisons that controls for type I errors. A P-value threshold of less than 0.05 was established as the criterion for statistical significance.

## Acknowledgements

We acknowledge Qiang Guo (State Key Laboratory of Oral Diseases, Sichuan University) for his excellent assistance in the μCT experiment. We thank Ning Ji, Xuelin Huang, and Yuwen Luo (Animal Care Center of State Key Laboratory of Oral Diseases, Sichuan University) for animal care and Mi Su (Functional Laboratory at the West China School of Basic Medical Sciences & Forensic Medicine) for the technical support. This work was supported by National Natural Science Foundation of China (82001062, 11932014, 82222015, and 82171001), China Postdoctoral Science Foundation (2022M722249), Research Funding from West China School/Hospital of Stomatology Sichuan University (RCDWJS2023-1). and National Key Research and Development Program of China (2023YFC2413600).

## Additional information

### Author contributions

Conceptualization: DZ, CZ. Methodology: ZW, QC, DZ. Investigation: YZ, ML, QG. Visualization: LZ, JW, YS. Funding acquisition: XL, DZ, CZ. Project administration: JJ, JX, XL. Supervision: XL, SZ. Writing – original draft: ZW, DZ. Writing – review & editing: ZW, DZ, CZ.

### Ethics

All animal procedures were approved by the Animal Care and Use Committee of Sichuan University (WCHSIRB-D-2023-288).

## Supplementary Materials

**Figure S1.**
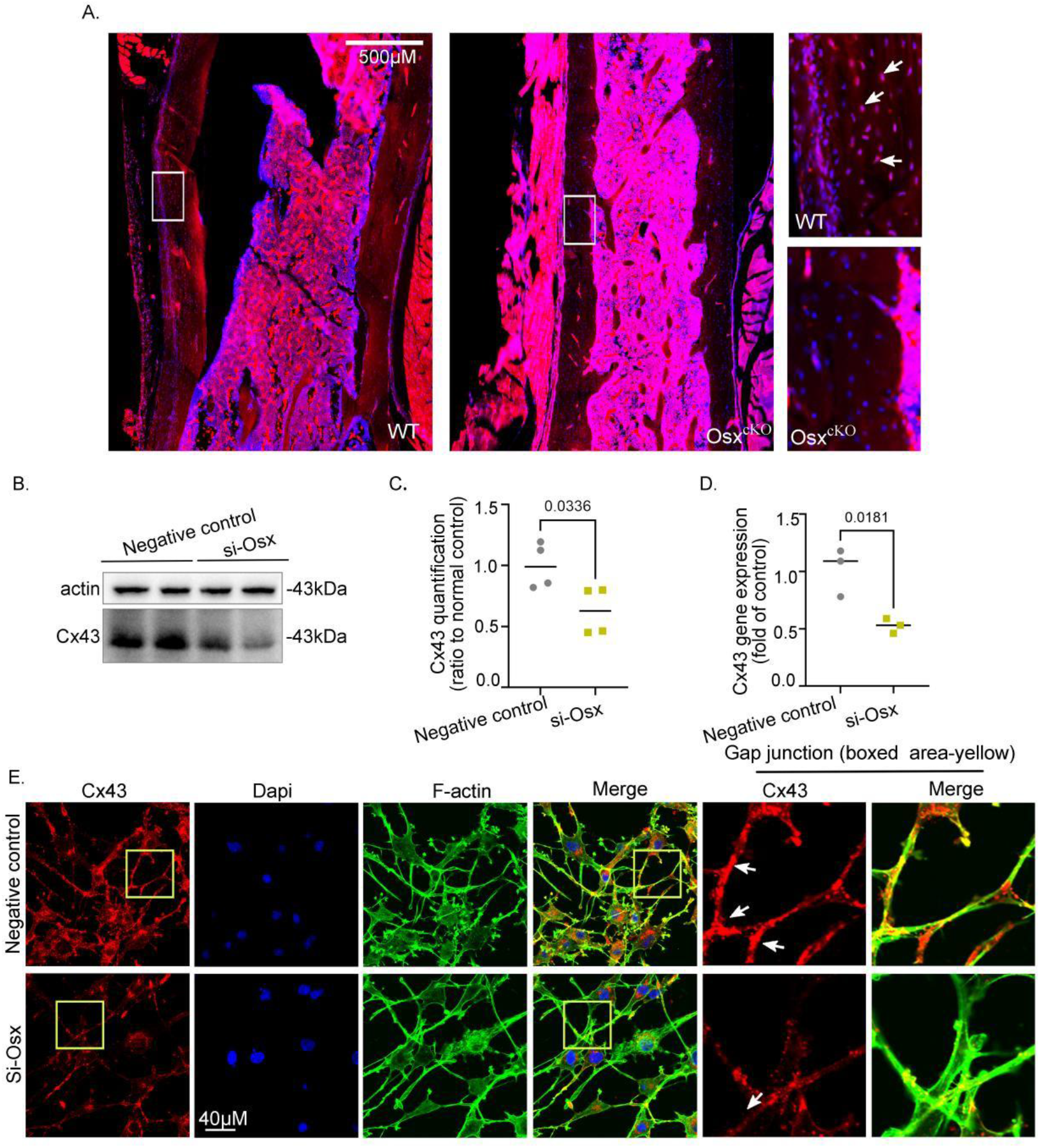
**The expression of Cx43 was reduced in Osx-deficient osteocytes both in vitro and in vivo.** (A) Immunofluorescence staining substantiated the decreased expression of Cx43 in the Osx^cKO^ mice. Scale bar, 500 μM. (B and C) Western blots revealed a reduction in Cx43 expression in Mlo-y4 osteocytes transfected with si-Osx. n = 3, with statistical analysis normalized to the control group. * p < 0.05. (D) qPCR showed the mRNA levels of Cx43 decreased in si-Osx Mlo-y4 osteocytes. n = 3. *p < 0.05. (E) Immunofluorescence staining showed a reduced expression of Cx43 in Mlo-y4 osteocytes with si-Osx as compared to the control group. The cytoskeleton was marked in green, Cx43 in red, and nuclei in blue. n = 3. Scale bar, 40 μM.

**Table S1.**
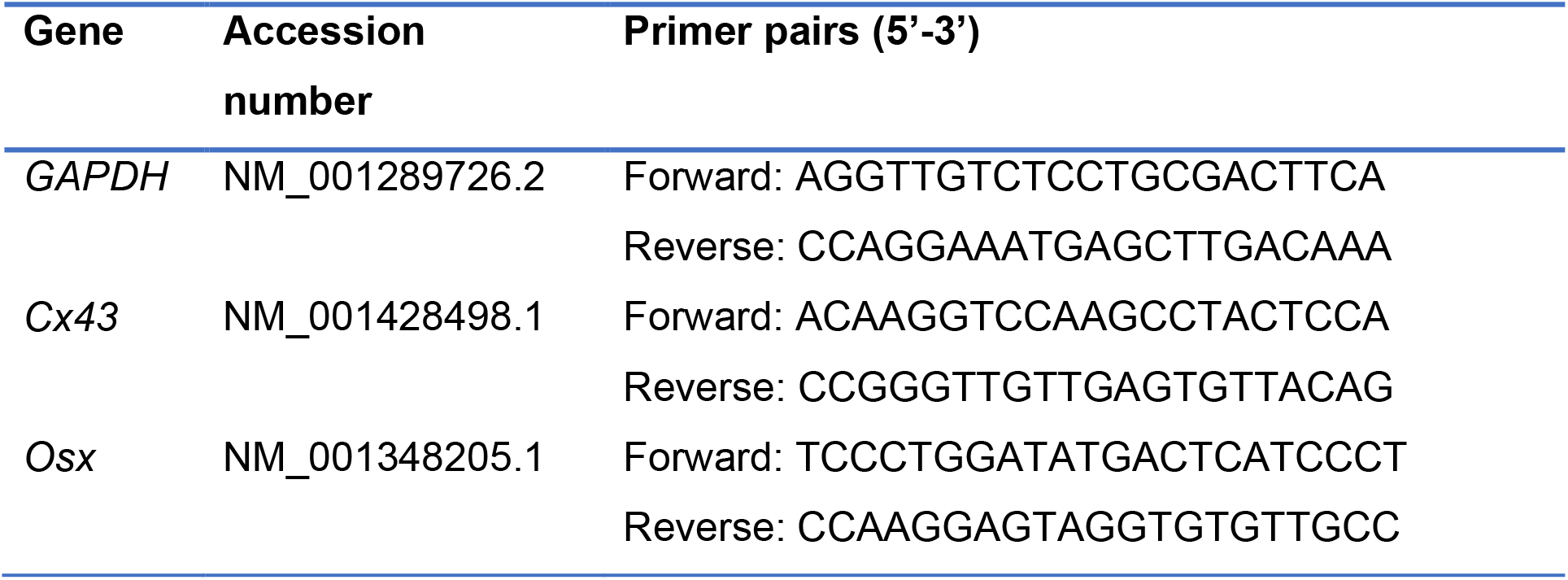
Sequences of primers for qPCR.

